# Symbiosis with rhizobia limits range expansion only in polyploid legumes

**DOI:** 10.1101/2022.03.01.482489

**Authors:** Zoe A. Parshuram, Tia L. Harrison, Anna K. Simonsen, John R. Stinchcombe, Megan E. Frederickson

## Abstract

- Both mutualism and polyploidy are thought to influence invasion success in plants but few studies have tested their joint effects. Mutualism can limit range expansion when plants cannot find a compatible partner in a novel habitat, or facilitate range expansion when mutualism increases a plant’s niche breadth. Polyploids are also expected to have greater niche breadth because of greater self-compatibility and phenotypic plasticity, increasing invasion success.
- For 839 legume species, we compiled data from published sources to estimate ploidy, symbiotic status with rhizobia, specificity on rhizobia, and the number of introduced ranges.
- We found that diploid species have had limited spread around the globe regardless of whether they are symbiotic or how many partners of rhizobia they can host. Polyploids, in contrast, have been successfully introduced to many new ranges, but interactions with rhizobia constrain their range expansion. In a hidden state model of trait evolution, we also found evidence of a high rate of re-diploidization in symbiotic legume lineages, suggesting that symbiosis and ploidy may interact at macroevolutionary scales.
- Overall, our results suggest that symbiosis with rhizobia affects range expansion only in polyploid legumes.

## Introduction

In plants, both polyploidy and mutualism are thought to contribute to ecological and evolutionary success. Polyploidization is an important driver of plant speciation, because it confers instant reproductive isolation and is associated with niche and range expansion (Soltis & Soltis, 2016; López-Jurado *et al*., 2019; Sheth *et al*., 2020). Likewise, mutualism has been linked with high rates of lineage diversification and is thought to increase ecological opportunity by expanding niche breadth and giving rise to coevolution with mutualist partners (Gómez & Verdú, 2012; Hembry *et al*., 2014; Weber & Agrawal, 2014; Zeng & Wiens, 2021). However, some work has also shown that engaging in mutualism can slow diversification (Kaur *et al*., 2019) and that relying on a mutualistic partner can limit range expansion (Simonsen *et al*., 2017). Despite the importance of both polyploidy and mutualism in determining where plants establish and persist, we do not know whether or how these factors interact to shape plant geographic ranges (Segraves & Anneberg, 2016). Here, we test whether ploidy and symbiosis jointly impact invasion success in legumes.

There are several reasons why polyploids may be better invaders than diploids. Polyploids generally have greater genetic variation (Otto & Whitton, 2000) and phenotypic plasticity (Mattingly & Hovick, 2021), both of which may allow polyploids to establish in and rapidly adapt to novel habitats. Polyploid plants also often have higher rates of self-fertilization, a trait that is associated with greater invasion success (Barringer & Geber, 2008). However, tests of polyploidy’s effects on range size have had mixed results. In the genus *Clarkia*, polyploids have larger ranges than diploids (Lowry & Lester, 2006), consistent with the above predictions. In contrast, in the Potentilleae tribe of Rosaceae, polyploids have smaller range sizes than diploids (Brittingham *et al*., 2018). Because species with higher ploidy tend to occupy more extreme habitats, they could become niche specialists, constraining their spread to new environments (Hummer, 2012; López-Jurado *et al*., 2019). Although legumes vary markedly in ploidy, no study has tested whether polyploid legumes have spread to more ranges than diploid legumes.

Legumes rely on bacteria partners called rhizobia for survival when living in nutrient-poor soil. Rhizobia form nodules on plant roots where rhizobia fix atmospheric nitrogen into a readily available form of nitrogen for plants (van Rhijn & Vanderleyden, 1995). The legume-rhizobium symbiosis is over 100 million years old but nonetheless not all legumes nodulate. It is unclear if nodulation has evolved multiple times after an old predisposition event (Doyle, 2011; Werner *et al*., 2014) or if the symbiotic trait has a single evolutionary origin followed by multiple losses across the legume phylogeny (Griesmann *et al*., 2018).

Although mutualism can facilitate range expansion in some plant species (Afkhami *et al*., 2014), a previous global analysis of legumes showed that symbiosis with rhizobia limits range expansion (Simonsen *et al*., 2017). Legumes that depend on rhizobia for nitrogen may be unable to establish in a new range if they cannot find compatible symbionts there. Expanding on the analysis of Simonsen *et al*. (2017), Harrison *et al*. (2018) found that symbiotic legumes that associate with many rhizobia taxa have spread to more new ranges than symbiotic legumes that specialize on just one or a few rhizobia partners, again suggesting that the availability of compatible rhizobia constrains the spread of legumes around the globe. However, neither of these previous studies considered ploidy.

Ploidy is predicted to have important effects on the interaction between legumes and rhizobia. Autotetraploid plants obtain more fixed nitrogen from larger nodules compared to diploids (Forrester & Ashman, 2020). Polyploid plants can also be more generalized on rhizobia, obtaining greater benefits from a wider diversity of rhizobia partners compared to diploids (Forrester *et al*., 2020). Legumes vary substantially in ploidy and polyploidy influences plant interactions with other species (Segraves & Anneberg, 2016), yet we currently lack broad-scale studies of how ploidy and symbiosis with rhizobia jointly influence invasion success in legumes. In this study, we 1) ask whether ploidy and symbiosis interact to affect range expansion in legumes and 2) estimate transition rates in symbiotic status and ploidy across the legume phylogeny to better understand how the symbiosis with rhizobia evolved in plants (e.g., whether it depends on evolutionary transitions in ploidy).

## Materials and Methods

We used published data to assemble a global trait dataset that included information on ploidy, mutualists, and geographic ranges for a total of 839 legume species.

### Ploidy

We estimated ploidy (i.e., the number of copies of each chromosome in a cell) for 839 species of legumes using methods adapted from Brittingham *et al*. (2018). Specifically, we extracted total chromosome count values for each species from the Chromosome Count Database (CCDB; Rice *et al*., 2015), the Index to Plant Chromosome Numbers (IPCN; Goldblatt & Johnson, 1979), or Rice *et al*. (2019). If a species was not found in one of these databases, we searched Web of Science for a genus-level average for the total chromosome count number (Supp Table 1). When multiple sources reported different chromosome counts for the same species, we used the median value in our analyses. To calculate ploidy for each species, we divided the chromosome counts by the genus-level base chromosome number as reported in Bairiganjan & Patnaik (1989) and Federov (1969). If the base chromosome number for a genus was missing from the Federov (1969) table, chromosome counts were divided by the base chromosome number reported for the legume subfamily: Mimosoideae (x= 13; Santos *et al*., 2012), Caesalpiniaceae (x = 7; Resende *et al*., 2013), or an average value for Papilionaceae (x = 10.5; Lackey, 1980) to find the ploidy level of the species. We categorized all species with ploidy values equal to or less than two as diploids and all species with ploidy values greater than two as polyploids. Although subfamily base chromosome number and genus base chromosome number values were significantly correlated in a Kendall’s rank test (tau=0.413, p<0.0001), we expect the ploidy values calculated from subfamily base chromosome numbers to be imperfect. However, the dataset with ploidy calculated using only the genus-level base chromosome values had fewer species overall (n=664) and only two species in it are non-symbiotic polyploids (Supp Table 2). These two species, *Cassia fistula* and *C. grandis*, are both ornamental trees and have a relatively high number of human uses in our dataset (7 uses and 4 uses, respectively) compared to a mean of 2.9 human uses for symbiotic species and a mean of 1.4 uses for non-symbiotic species. It is thus difficult to disentangle the influence of the number of human uses, ploidy, and lack of nodules on the number of introduced ranges in these two species. Therefore, for our analyses we used a combination of genus-level and subfamily-level estimated ploidy levels (n=839). Specifically, if ploidy calculated from the genus-level base chromosome number was available, we used that value, and if it was not available, we used ploidy calculated using the subfamily-level base chromosome number. Since there was a small number of polyploid species calculated from subfamily base chromosome values, we searched the literature to confirm that these species are indeed polyploids (Supp Table 3). Of these 62 species, 28 were confirmed to be polyploids, 22 were changed to diploid in our dataset, and 12 were left as polyploid because we could not find any information on ploidy level. We then chose 62 species at random from the list of diploids in the dataset and looked in the literature to confirm their ploidy level. Of these 62 species, 2 were changed to polyploid, 14 did not have ploidy levels reported in the literature, and the rest were confirmed to be diploid (Supp Table 4), suggesting that our method incorrectly classified diploids in only ~3% of species. After these corrections, our dataset consisted mostly of diploid species (n = 568) and fewer polyploid species (n = 271; Supp Table 2).

### Mutualists

We obtained data on the symbiotic status of the legumes in our study from Werner *et al*. (2014). If a species is known to form nodules with rhizobia, it was categorized as a symbiotic species. We used the dataset assembled by Harrison *et al*. (2018) to determine whether each symbiotic legume species is a specialist or a generalist with regards to its interactions with rhizobia. Harrison *et al*. (2018) used data from Andrews and Andrews (2017) to determine the number of unique rhizobia genera that interact with diverse legume species. We classified a legume species as a specialist if it interacts with only one genus of rhizobia. Any legume species that interacts with more than one genus of rhizobia is a generalist.

### Legume ranges

We obtained data on the number of introduced ranges for each legume species from Simonsen *et al*. (2017). Using geographic data from the International Legume Database and Information Service (ILDIS), Simonsen *et al*. (2017) classified native and invaded ranges for each species as discrete geographical ranges that roughly align with geopolitical boundaries. If a species was present in a non-native region and the neighbouring regions, it was counted as a successful invasion event, or one introduced range. We also included in our models several other covariates from the Simonsen *et al*. (2017) dataset, specifically the number of human uses for each legume species, the area of a species’ native range, the midpoint latitude of a species’ native range, and whether or not the species was annual or perennial.

### Statistical analyses

All analyses were performed in R (R Core Team, 2020). We used the *gls* function in the package *nlme* (Pinheiro *et al*., 2020) to fit phylogenetic least-squares models (PGLS). We pruned the Zanne *et al*. (2014) angiosperm phylogeny to retain only the species in our trait dataset. First, we modeled the number of introduced ranges as a function of the main and interactive effects of symbiotic status (symbiotic/non-symbiotic) and ploidy (diploid/polyploid). However, because the trait dataset is highly unbalanced (Supp Table 2), we also split up this dataset into symbiotic species and non-symbiotic species to analyze the impact of ploidy on range expansion separately for these two categories of legumes. For only the symbiotic species in the dataset, we also modeled the main and interactive effects of specialization (specialist/generalist) and ploidy (diploid/polyploid) on introduction success. We treated symbiotic status, specialization, and ploidy as binary, categorical variables. All analyses included the following covariates: number of human uses, scaled value of total native area, life cycle duration (annual, perennial, etc.), and absolute latitude of origin. We allowed Pagel’s lambda parameter to vary and optimize in the PGLS models by setting the fixed argument to false. The branch lengths of the phylogeny are multiplied by this lambda parameter to account for phylogenetic signal in the data. A lambda value of 0 indicates that variation in the data is independent of phylogeny and a lambda value of 1 indicates a Brownian motion model of evolution. We logged (base 10) the number of introduced ranges (plus one to avoid zero values) in order to improve normality and homoscedasticity. When the PGLS models estimated negative lambda values (suggesting phylogenetic overdispersion in the data) or values very close to zero (suggesting very weak phylogenetic structure in the data) we also performed tests on the data using the *glm* function without accounting for phylogeny. Since the number of introduced ranges was not normally distributed and overdispersed, we fit the data to a quasi-poisson distribution. We tested for significance by performing a type III (on models with interaction terms) or a type II (on models without interaction terms) ANOVA using the *Anova* function in the *car* package (Fox & Weisberg, 2019).

### State transitions

We modeled the evolution of ploidy and symbiosis across the Zanne *et al*. (2014) phylogeny with the package *corHMM* (Beaulieu *et al*., 2021) to estimate transitions between the following states: non-symbiotic diploid, non-symbiotic polyploid, symbiotic diploid, and symbiotic polyploid. We fit a simple corHMM model with no hidden states (i.e., a single rate category) and also a model with hidden states (i.e., two rate categories) on the genus/subfamily-level ploidy data. Because the hidden state model always outperformed the model with no hidden states, we plotted transition rates estimated from the hidden state models on the phylogeny using the plotSimmap function in *phytools* (Revell, 2012).

## Results

### Symbiotic status and ploidy

Non-symbiotic polyploids have been successfully introduced to more new ranges than symbiotic polyploids (Fig 1a). In contrast, diploid species have been introduced to few new ranges regardless of symbiotic status (Fig 1a). Symbiotic status and ploidy interacted significantly to predict legume introduced ranges, although only in a non-phylogenetic model (Table 1). In the PGLS model that accounts for phylogeny, there was a non-significant interaction effect between symbiotic status and ploidy on the number of introduced ranges (Supp Table 5). However, the PGLS model also showed a weak phylogenetic signal in the number of introduced ranges (λ= 0.0431). Furthermore, when we analyzed ploidy’s effects separately within non-symbiotic and symbiotic legumes, we found a significant effect of ploidy on range expansion only in non-symbiotic legumes (with moderate lambda value of λ= 0.3552), but not in symbiotic legumes (Supp Table 6). In our second set of analyses on symbiotic species only, polyploid generalists were introduced to many more ranges than diploid generalists and both diploid and polyploid specialists (Fig 1b). There was a significant interaction between specialization on rhizobia and ploidy in both our phylogenetically corrected (Supp Table 5) and uncorrected models (Table 1).

**Figure 1.**
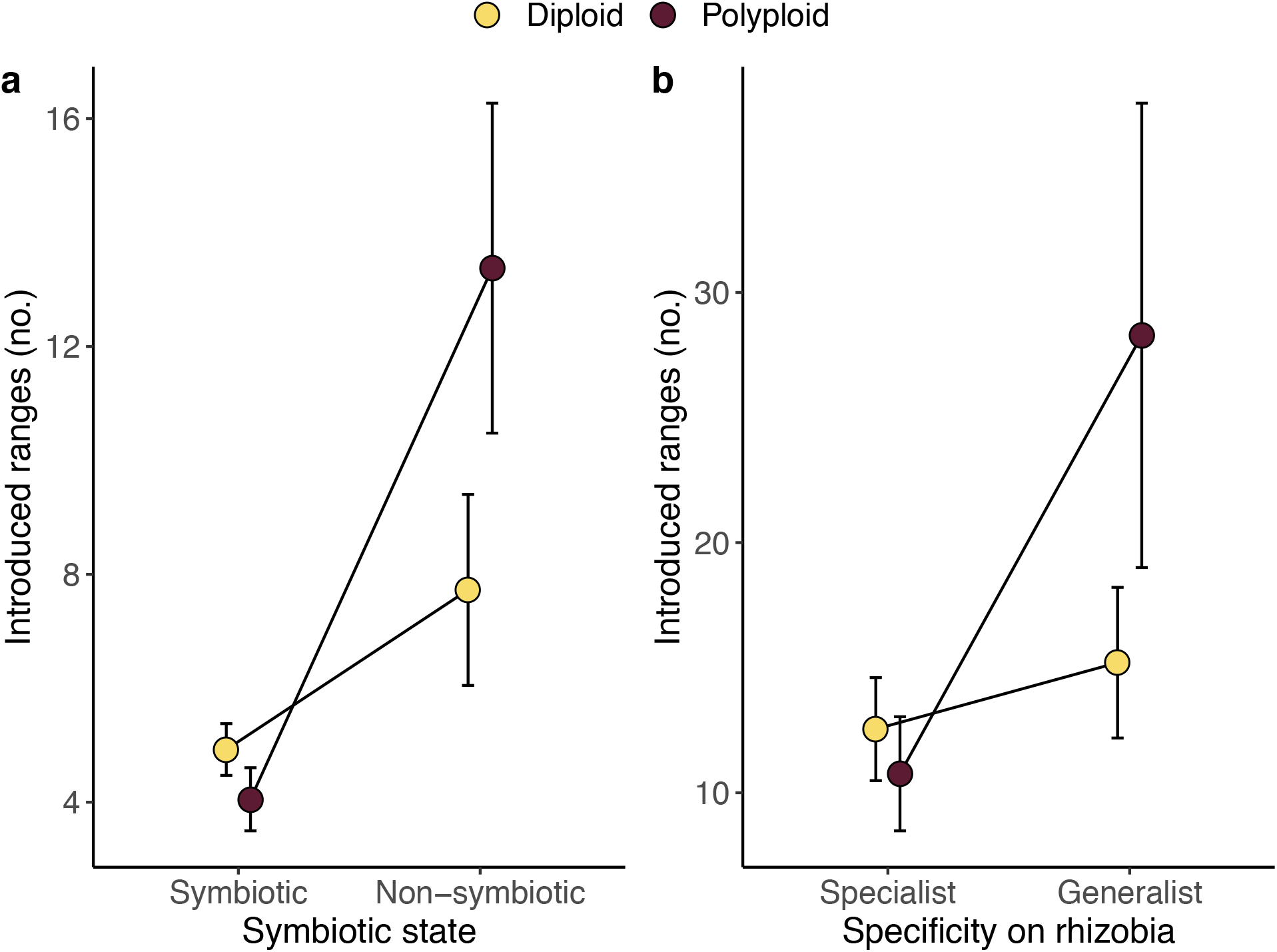
Mean (±1SE) number of introduced ranges for diploid and polyploid legumes that a) form nodules with rhizobia (‘symbiotic’) or do not form nodules (‘non-symbiotic’) and b) associate with only one genus of rhizobia (‘specialist’) or associate with more than one genus of rhizobia (‘generalist’).

**Table 1.** Estimates and results for effects of ploidy and symbiotic status on the number of introduced ranges in legumes obtained from glm models fit with a quasi-poisson distribution. Ploidy is estimated from combined genus and subfamily-level base chromosome numbers. Italicized factors are significant at p < 0.05 and p values reported here are the results of type III ANOVAs. Bolded factors highlight significant interaction effects on legume introductions. Ploidy was coded as 0=diploid and 1=polyploid while symbiotic status was coded as 0=non-symbiotic and 1=symbiotic in the model. Therefore, the intercept represents non-symbiotic diploids. Specialists were coded as 1 and generalists as 0 in the model.

| Factor | Estimate | s.e. | d.f. | Wald $\chi^2$ | p value |
| --- | --- | --- | --- | --- | --- |
| **Symbiosis status, d.f. = 810** |  |  |  |  |  |
| *Ploidy (polyploid)* | 0.573 | 0.210 | 1 | 6.530 | 0.0106 |
| Symbiosis (symbiotic) | 0.248 | 0.178 | 1 | 2.300 | 0.1295 |
| *Total native area* | -0.121 | 0.040 | 1 | 13.730 | 0.0002 |
| *Human uses* | 0.432 | 0.018 | 1 | 595.820 | < 0.0001 |
| *Absolute latitude* | -0.011 | 0.004 | 1 | 12.160 | 0.0005 |
| *Annual* | 0.405 | 0.137 | 1 | 11.940 | 0.0005 |
| ***Ploidy\*Symbiosis*** | **-0.649** | **0.243** | **1** | **6.860** | **0.0088** |
| **Specialist status, d.f. = 129** |  |  |  |  |  |
| Ploidy (polyploid) | 0.498 | 0.252 | 1 | 3.114 | 0.0776 |
| Specificity (specialist) | 0.116 | 0.178 | 1 | 0.420 | 0.5167 |
| *Total native area* | -0.220 | 0.044 | 1 | 22.078 | < 0.0001 |
| *Human uses* | 0.290 | 0.030 | 1 | 89.031 | < 0.0001 |
| *Absolute latitude* | -0.020 | 0.005 | 1 | 15.868 | < 0.0001 |
| Annual | 0.239 | 0.166 | 1 | 1.764 | 0.1841 |
| ***Ploidy\*Specificity*** | **-0.931** | **0.312** | **1** | **6.640** | **0.0099** |

### State Transitions

The hidden state model performed better (-lnL= −389.030, AIC=814.060, n=839) than the simple model (-lnL= −439.715, AIC=895.603, n=839) therefore we present results only for the hidden state model. The highest transition rate (100.00) was from polyploidy to diploidy within symbiotic lineages in the first rate class (Table 2). Overall, transitions in ploidy were higher within symbiotic lineages than in non-symbiotic lineages. Within non-symbiotic lineages, transitions to diploidy were higher (5.0949) than transitions to polyploidy (1.6291).

**Table 2.**
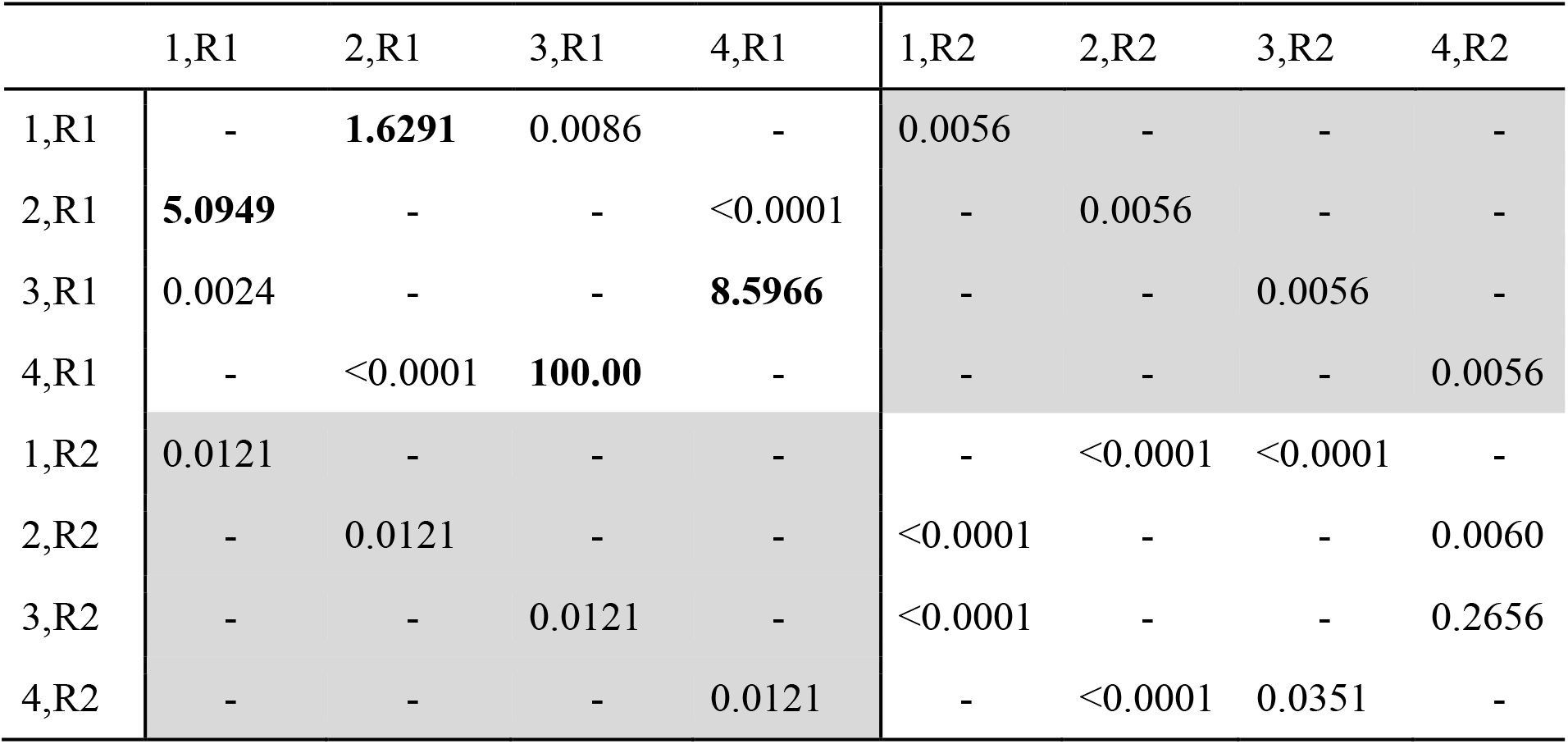
corHMM results for hidden state model for the genus and subfamily level ploidy values. The white quadrants indicate observed state transitions within rate categories and grey quadrants indicate state transitions between rate categories. The different states are represented by 1=non-symbiotic diploid, 2=non-symbiotic polyploid, 3=symbiotic diploid, 4=symbiotic polyploid. The two rate classes are represented as R1 and R2. Highest transition rates are in bold. Transitions are read from row to column.

|  | 1,R1 | 2,R1 | 3,R1 | 4,R1 | 1,R2 | 2,R2 | 3,R2 | 4,R2 |
| --- | --- | --- | --- | --- | --- | --- | --- | --- |
| 1,R1 | - | **1.6291** | 0.0086 | - | 0.0056 | - | - | - |
| 2,R1 | **5.0949** | - | - | <0.0001 | - | 0.0056 | - | - |
| 3,R1 | 0.0024 | - | - | **8.5966** | - | - | 0.0056 | - |
| 4,R1 | - | <0.0001 | **100.00** | - | - | - | - | 0.0056 |
| 1,R2 | 0.0121 | - | - | - | - | <0.0001 | <0.0001 | - |
| 2,R2 | - | 0.0121 | - | - | <0.0001 | - | - | 0.0060 |
| 3,R2 | - | - | 0.0121 | - | <0.0001 | - | - | 0.2656 |
| 4,R2 | - | - | - | 0.0121 | - | <0.0001 | 0.0351 | - |

## Discussion

### Symbiosis and ploidy interact to affect range expansion

Overall, we found that symbiosis only impacts range expansion within polyploid species and that diploid legumes spread to few new ranges regardless of symbiotic status. Therefore, our results generally support the hypothesis that polyploid legumes are niche generalists and better able to colonize new habitats. However, when polyploids are also symbiotic they are restricted in their range expansion, suggesting that symbiotic legumes have difficulty finding a compatible rhizobia partner when they are introduced to a novel habitat (Simonsen *et al*., 2017). Nonetheless, in legumes, diploidy seems to be the main factor restricting a plant’s range; both non-symbiotic diploids and generalist diploids were introduced to few new ranges despite not requiring or not specializing on rhizobia.

Polyploids might be good colonizers because they have more genetic material in their large genomes giving them greater adaptive potential (Otto & Whitton, 2000). Polyploids also often have greater capacity for phenotypic plasticity and can be better competitors than diploids, since they have fast germination and growth rates (te Beest *et al*., 2012). However, being unable to associate with compatible rhizobia seems to cancel out the advantages of being polyploid. One explanation is that these symbiotic polyploids are largely specialists in terms of which rhizobia species they can form nodules with, making it more challenging to find beneficial rhizobia in new habitats. There were more specialist polyploids (25 species) than generalists (7 species) in our dataset, in contrast to previous results that report generalization on rhizobia in polyploids (Forrester *et al*., 2020). In our analyses on symbiotic species only, we found that generalist polyploids were introduced to far more ranges than any other category of legume (Fig 1b), providing further support for the theory that having many potential rhizobia partners is beneficial for range expansion, but only when paired with the benefits of being a polyploid.

Some caveats are in order, however. There were somewhat conflicting results for the interaction between symbiosis and ploidy in our PGLS analysis (non-significant) and non-phylogenetic GLM (significant). However, the PGLS model has less statistical power because it includes a phylogenetic correction, perhaps explaining the discrepancy, and the phylogenetic signal was very low in the PGLS model so phylogenetic correction may be unnecessary. We also used a combination of ploidy estimates calculated from genus- and subfamily-level base chromosome values. Although this allowed us to compile a larger dataset, it also introduced some error in ploidy estimates since subfamily base values are less reliable. The species for which we have more accurate ploidy estimates (i.e., genus-level data) are likely well-studied species that have spread widely across the globe. We corrected the data for some taxa initially categorized as polyploids, and a smaller number initially categorized as diploids, by searching the literature to help deal with this issue.

### High rate of re-diploidization in legumes

Overall, the best-fitting model of trait evolution was a hidden state model, likely because other unmeasured traits contribute to rate heterogeneity across such a large and old clade as the legumes (Beaulieu & O’Meara, 2016). This model found that evolutionary transitions in ploidy were generally higher within symbiotic than non-symbiotic lineages. Within symbiotic lineages, we observed an especially high rate of re-diploidization. Re-diploidization is a common process that occurs in many plant lineages (Tamayo-Ordóñez *et al*., 2016) and seems to occur more frequently in symbiotic than non-symbiotic legume lineages. Polyploid species may undergo re-diploidization if the extra genetic material in the genome causes dosage imbalance, errors in mitosis or meiosis, disruption to gene regulation, or epigenetic instability (Comai, 2005). Larger genomes are also generally more costly to maintain. After polyploid legumes evolved genes important for interactions with rhizobia, they may have experienced extensive gene loss (and thus genome reduction) during re-diploidization which would reduce these costs.

The ancestral state in the legume tree was non-symbiotic and diploid. Our analysis suggests that polyploidy directly evolved from diploidy, leading to the evolution of symbiosis followed by re-diploidization in symbiotic species (Fig. 2). There is evidence in the family Brassicaceae that whole genome duplication has facilitated the evolution of diverse chemical defense compounds against herbivores (Edger *et al*., 2015). If polyploidy did indeed evolve before symbiosis with rhizobia, the extra genetic material in the polyploid genome could provide more opportunity for mutations and the evolution of symbiosis to occur (Soltis & Soltis, 2016). However, previous in-depth analysis of transcriptome data in a few key legume species suggests that the evolution of polyploidy and symbiosis in legumes are unrelated (Cannon *et al*., 2010). Overall, our analysis still suggests that polyploidy evolved first and is a potential predisposition event to the evolution of symbiosis. However, this is an important question that warrants further exploration with a larger dataset and better ploidy estimates.

**Figure 2.**
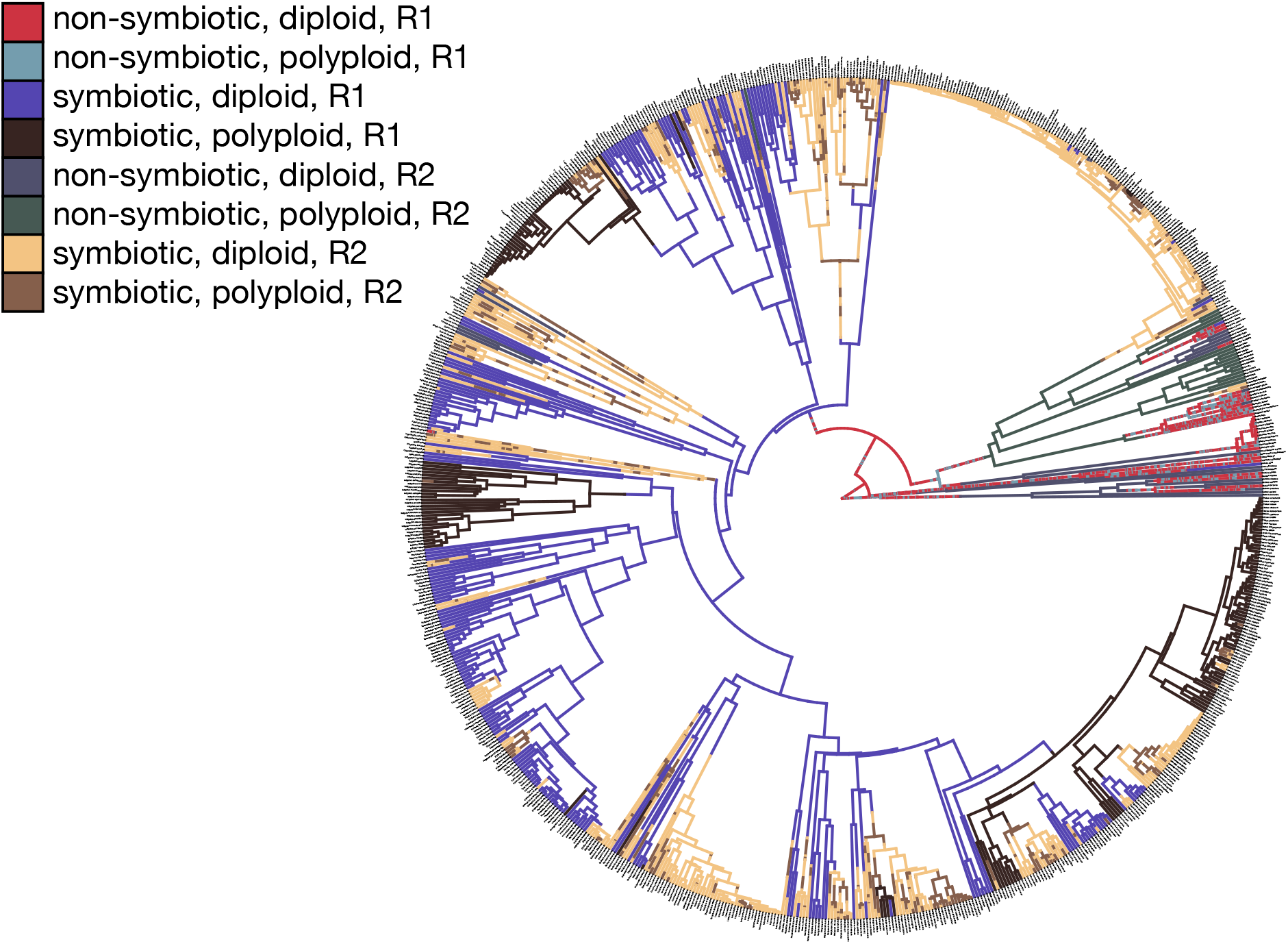
Transition rates between four states (‘non-symbiotic diploid’, ‘non-symbiotic polyploid’, ‘symbiotic diploid’, ‘non-symbiotic polyploid’) in two rate classes (R1 and R2) estimated from a corHMM (hidden state) model. Ploidy is estimated from combined genus and subfamily-level base chromosome numbers.

### Conclusion

Our results show the importance of considering the effects of both ploidy and symbiosis on invasion success; here, we showed that ploidy can influence the impact mutualism has on the spread of invasive species across the globe. Previous work has shown that symbiosis (Simonsen *et al*., 2017) and specialization on a small number of rhizobia partners (Harrison *et al*., 2018) introduces a barrier to range expansion in legumes. Our study supports these results but shows that this barrier only affects polyploid legumes which otherwise have an advantage over diploids in terms of successful introductions to new ranges. Looking ahead, the joint influence of ploidy and mutualism on invasion success should also be evaluated in other mutualisms, such as plant-pollinator interactions; polyploids tend to be self-fertilizing, which is thought to facilitate range expansion (Barringer & Geber, 2008), but some work has suggested that invasive plants can take advantage of local pollinators (Graves & Shapiro, 2003). Our results also suggest that evolutionary transitions in ploidy and symbiosis might be linked at macro-evolutionary scales; polyploidy appears to have arisen before symbiosis in legumes, perhaps setting the stage for the evolution of the legume-rhizobium mutualism. Nonetheless, once symbiotic, legumes frequently revert to their ancestral diploid state, likely explaining why so many extant symbiotic legumes are diploids.

## Acknowledgements

We would like to thank Nikki Forrester for her methodological advice on estimating ploidy levels in our global legume dataset. Our work was funded by an NSERC URSA (ZAP), an NSERC PGS D scholarship and QEII-GSST award (TLH), and an NSERC Discovery Grant (MEF).

## Data Availability

All data and code for reproducing our analyses will be made available on GitHub at the time of publication.

