## Supplemental Data 1 for "Symbiosis with rhizobia limits range expansion only in polyploid legumes"

**Supplemental file**

These are the tables and supplemental material to accompany the main text of the manuscript titled "Symbiosis with rhizobia limits range expansion only in polyploid legumes".

**Supplemental Table 1.** Base chromosome counts for different legume genera and subfamilies obtained from the literature.

| Genus & Base Chromosome Count | Reference |
|---|---|
| Acacia = 26 | Blakesley D, Allen A, Pellney TK and Roberts AV. 2002. Natural and Induced Polyploidy in *Acacia dealbata Link. and Acacia mangium Willd*. *Annals of Botany* 90: 391-398. |
| Calliandra = 16,<br>Mimosa = 26,<br>Piptadenia = 26 | Santos CXC, Carvalho R, Almeido EM, and Felix LP. 2012. Chromosome number variation and evolution in Neotropical Leguminoseae (Mimosoideae) from northeastern Brazil. *Genetics and Molecular Research* 11: 2451-75. |
| Alhagi = 16 | Shedai M, Yazdanbakhsh Z, Assadi M, Moussavi M. 2008. Cytology and morphometry study of Alhagi (Leguminosae) species in Iran. *Nordic Journal of Botany* 21: 83-91. |
| Centrolobium = 20 | Dahmer N, Wittman MTS, Kaminski PE. 2009. Chromosome number and karyotype of the endangered Amazonian woody *Centrolobium paraense Tul*. Species. *Crop Breeding and Applied Biotechnology* 9: 382-385. |
| Hypocalyptus = 20 | Schutte AL, van Wyk BE. 1998. The Tribal Position of Hypocalyptus Thunberg (Fabaceae). *Novon* 8: 178-182. |
| Lablab = 23 | Shivashankar G, Kulkarni RS. 1989. *Lablab purpureus* L. (Sweet)In: van der Maesen LJG, Somaatmadja S, editors. Plant Resources of South-East Asia No. 1: Pulses. Wageningen, The Netherlands: Pudoc 48-50. |
| Macroptilium = 22 | Rojas-Sandoval J. 2018. *Macroptilium atropurpureum* (siratro). *Invasive Species Compendium*. Wallingford, UK: CABI. doi:10.1079/ISC.110272.20203483153 |

| | |
|---|---|
| Macrotyloma = 20 | Halder S, Datta AK, Mandal A, Ghosh BK. 2012. *Macrotyloma uniflorum* (Lam.) Verdc. (Leguminosae)-A Note on Chromosomal Studies. *Cytologia.* doi: 10.1508/cytologia.77.447 |
| Cystisus = 46 | Maude PF. 2006. Chromosome numbers in some British Plants. *New Phytologist.* doi: 10.1111/j.1469-8137.1940.tb07119.x |
| Ammopiptanthus = 18 | Zhang S-Z, Cao R. 1990. Study on Chromosome Number and Karyotype of *Ammopiptanthus mongolicus*[J]. *Journal of Systematics and Evolution* 28: 133-135. |
| Archidendron = 26 | PlantUse English contributors. 2016. *Archidendron (PROSEA)*. *PlantUse English.* [WWW document] URL https://uses.plantnet-project.org/e/index.php?title=Archidendron (PROSEA)&oldid=210426. [accessed July 5, 2021]. |
| Bossiaea = 16 | Ross, J H. 2006. A Conspectus of the Western Australian Bossiaea species (Bossiaeeae: Fabaceae). *Muelleria 23: 15-143.* |
| Brachystegia = 24 | Watson L, Dallwitz MJ. 1993. Delta-inkey.com. The genera of Leguminosae-Caesalpinioideae and Swartzieae: descriptions, illustrations, identification, and information retrieval. [WWW document] https://www.delta-intkey.com/caes/en/www/brachyst.htm. [accessed July 5, 2021]. |
| Senna = 28 | Marques de Resende KF, Davide LC, Torres GA. 2013. Chromosome number and meiosis in populations of *Senna* species (Caesalpinioideae – Fabaceae) from Southeast Brazil. *Caryologia* doi: 10.1080/00087114.2012.760883 |

**Supplementary Table 2.** Summary of the number of plant species in each category of data. Ploidy levels in the genus/subfamily dataset have been corrected for ploidy reported in the literature.

| Ploidy | Symbiosis status | Number of species |
|---|---|---|
| **Genus-level ploidy dataset** | | |
| Diploid | Non-symbiotic | 21 |
| Diploid | Symbiotic | 412 |
| Polyploid | Non-symbiotic | 2 |
| Polyploid | Symbiotic | 229 |
| **Genus/subfamily level ploidy dataset** | | |
| Diploid | Non-symbiotic | 44 |
| Diploid | Symbiotic | 523 |
| Polyploid | Non-symbiotic | 24 |
| Polyploid | Symbiotic | 248 |

**Supplemental Table 3.** Polyploid species identified in our dataset. We looked up these species in the literature to confirm if they were polyploid or whether we should correct our dataset and change certain species to diploid.

| Species | Ploidy level in our dataset | Ploidy level in the literature | Reference |
|---|---|---|---|
| *Acrocarpus fraxinifolius* | polyploid | Not reported | NA |
| *Apuleia leiocarpa* | polyploid | diploid | Auler NMF and Battistin A (1999). Análise do cariótipo de Apuleia leiocarpa (Vog.) Macbr. Cienc. Rural 29: 167-169. |
| *Colvillea racemosa* | polyploid | Not reported | NA |
| *Copaifera langsdorffii* | polyploid | diploid | Daviña JR et al (2001) Chromosome studies on plants from Paraguay, International Journal of Experimental Botany, 215-224. |
| *Copaifera officinalis* | polyploid | likely polyploid | Al-Ahmad H, Kania SA, Trent DJ, Stewart CN (2017) Estimation of Nuclear DNA Contents of Three Economically Important Plant Species by Laser Flow Cytometry, An - Najah Univ. J. Res. (N. Sc.) 31(1):36-54. |
| *Delonix elata* | polyploid | relative *Delonix regia* is polyploid | La Fleur, Angulus L, A Study of Normally Occurring Polyploidi in the Development of Delonix Regia (Bojer) Rafinesque (1953). ETD Collection for Fordham University. AAI10992926. |
| *Distemonanthus benthamianus* | polyploid | Not reported | NA |
| *Gleditsia amorphoides* | polyploid | diploid | Schnabel A, Wendel JF (1998) Cladistic biogeography of Gleditsia (Leguminosae) based on ndhF and rpl16 chloroplast gene sequences, American Journal of Botany, 85 (12): 1753-1765. |
| *Gleditsia japonica* | polyploid | diploid | Schnabel A, Wendel JF (1998) Cladistic biogeography of Gleditsia |

| | | | |
|---|---|---|---|
| | | | (Leguminosae) based on ndhF and rpl16 chloroplast gene sequences, American Journal of Botany, 85 (12): 1753-1765. |
| *Gleditsia triacanthos* | polyploid | diploid | Schnabel A, Wendel JF (1998) Cladistic biogeography of Gleditsia (Leguminosae) based on ndhF and rpl16 chloroplast gene sequences, American Journal of Botany, 85 (12): 1753-1765. |
| *Guibourtia coleosperma* | polyploid | diploid/polyploid | Tosso F et al (2016) Microsatellite development for the genus Guibourtia (Fabaceae, Caesalpinioideae) reveals diploid and polyploid species, Applications in Plant Sciences, 4(7): 1600029.<br>Serbin GM et al (2019) Karyological traits related to phylogenetic signal and environmental conditions within the Hymnenaea clade (Leguminosae, Detarioideae), Perspectives in Plant Ecology, Evolution and Systematics, 39:125462. |
| *Gymnocladus chinensis* | polyploid | diploid | Schnabel A, Wendel JF (1998) Cladistic biogeography of Gleditsia (Leguminosae) based on ndhF and rpl16 chloroplast gene sequences, American Journal of Botany, 85 (12): 1753-1765.<br>Cannon et al (2014) Multiple Polyploidy Events in the Early Radiation of Nodulating and Nonnodulating Legumes, Mol. Biol. Evol, 32(1): 193-210. |
| *Gymnocladus dioica* | polyploid | diploid | Schnabel A, Wendel JF (1998) Cladistic biogeography of Gleditsia (Leguminosae) based on ndhF and rpl16 chloroplast gene sequences, American Journal of Botany, 85 (12): 1753-1765.<br>Cannon et al (2014) Multiple Polyploidy Events in the Early Radiation of Nodulating and Nonnodulating Legumes, Mol. Biol. Evol, 32(1): 193-210. |
| *Haematoxylum* | polyploid | Not reported | NA |

| | | | |
|---|---|---|---|
| *campechianum* | | | |
| *Hoffmannseggia glauca* | polyploid | diploid | Zanín LA, C, MA (2001) EL CARIOTIPO DE HOFFMANNSEGGIA GLAUCA (FABACEAE), Darwiniana, 39 (1-2): 11-13. |
| *Hymenaea courbaril* | polyploid | diploid | Serbin GM et al (2019) Karyological traits related to phylogenetic signal and environmental conditions within the Hymnenaea clade (Leguminosae, Detarioideae), Perspectives in Plant Ecology, Evolution and Systematics, 39:125462. |
| *Hymenaea parvifolia* | polyploid | diploid | Serbin GM et al (2019) Karyological traits related to phylogenetic signal and environmental conditions within the Hymnenaea clade (Leguminosae, Detarioideae), Perspectives in Plant Ecology, Evolution and Systematics, 39:125462. |
| *Hymenaea verrucosa* | polyploid | diploid | Serbin GM et al (2019) Karyological traits related to phylogenetic signal and environmental conditions within the Hymnenaea clade (Leguminosae, Detarioideae), Perspectives in Plant Ecology, Evolution and Systematics, 39:125462. |
| *Hymenostegia afzelii* | polyploid | Not reported | NA |
| *Parkinsonia aculeata* | polyploid | likely polyploid | Siliwinska E, Pisarczyk I, Pawlik A, Galbraith D (2009) Measuring genome size of desert plants using dry seeds. Botany 87: 127-135. |
| *Peltophorum dubium* | polyploid | diploid | Singhal VK, Gill BS, Sidhu MS (1990) Cytological explorations of Indian woody legumes. Proc. Indian Acad. Sci (Plant Sci) 100(5): 319-331.<br>Van-Lume B, Souza G (2018) Cytomolecular analysis of species in the Peltophorum clade (Caesalpinioideae, |

| | | | |
|---|---|---|---|
| | | | Leguminosae), Brazilian Journal of Botany, 41:385-392. |
| *Peltophorum pterocarpum* | polyploid | diploid | Singhal VK, Gill BS, Sidhu MS (1990) Cytological explorations of Indian woody legumes. Proc. Indian Acad. Sci (Plant Sci) 100(5): 319-331.<br>Van-Lume B, Souza G (2018) Cytomolecular analysis of species in the Peltophorum clade (Caesalpinioideae, Leguminosae), Brazilian Journal of Botany, 41:385-392. |
| *Pickeringia montana* | polyploid | Not reported | NA |
| *Pterogyne nitens* | polyploid | Not reported | NA |
| *Schizolobium parahyba* | polyploid | likely polyploid | Biondo E, Miotto, STS, Schifino-Wittmann MT (2005) Números cromossômicos e implicações sistemáticas em espécies da subfamília Caesalpinioideae (Leguminosae) ocorrentes na região sul do Brasil, Revista Brasil. Bot. 28 (4): 797-808. |
| *Schotia brachypetala* | polyploid | Not reported | NA |
| *Senna alexandrina* | polyploid | polyploid | Pereira Cordeiro JM & Felix, LP. (2018) Intra- and interspecific karyotypic variations of the genus Senna Mill. (Fabaceae, Caesalpinioideae), Acta Bot. Bras 32 (01): 128-134. |
| *Senna candolleana* | polyploid | polyploid | Pereira Cordeiro JM & Felix, LP. (2018) Intra- and interspecific karyotypic variations of the genus Senna Mill. (Fabaceae, Caesalpinioideae), Acta Bot. Bras 32 (01): 128-134. |
| *Senna didymobotrya* | polyploid | polyploid | Pellerin RJ, Waminal NE, Kim HH (2019) FISH mapping of rDNA and telomeric repeats in 10 Senna species, Horticulture, Environment, and Biotechnology, 60: 253-260. |
| *Senna* | polyploid | polyploid | Pereira Cordeiro JM & Felix, LP. |

| | | | |
|---|---|---|---|
| *obtusifolia* | | | (2018) Intra- and interspecific karyotypic variations of the genus Senna Mill. (Fabaceae, Caesalpinioideae), Acta Bot. Bras 32 (01): 128-134. |
| *Senna occidentalis* | polyploid | polyploid | Pellerin RJ, Waminal NE, Kim HH (2019) FISH mapping of rDNA and telomeric repeats in 10 Senna species, Horticulture, Environment, and Biotechnology, 60: 253-260. |
| *Senna pendula* | polyploid | polyploid | Pereira Cordeiro JM & Felix, LP. (2018) Intra- and interspecific karyotypic variations of the genus Senna Mill. (Fabaceae, Caesalpinioideae), Acta Bot. Bras 32 (01): 128-134. |
| *Senna reticulata* | polyploid | polyploid | Pereira Cordeiro JM & Felix, LP. (2018) Intra- and interspecific karyotypic variations of the genus Senna Mill. (Fabaceae, Caesalpinioideae), Acta Bot. Bras 32 (01): 128-134. |
| *Senna surattensis* | polyploid | polyploid | Pereira Cordeiro JM & Felix, LP. (2018) Intra- and interspecific karyotypic variations of the genus Senna Mill. (Fabaceae, Caesalpinioideae), Acta Bot. Bras 32 (01): 128-134. |
| *Senna tora* | polyploid | polyploid | Pellerin RJ, Waminal NE, Kim HH (2019) FISH mapping of rDNA and telomeric repeats in 10 Senna species, Horticulture, Environment, and Biotechnology, 60: 253-260. |
| *Amicia zygomeris* | polyploid | Not reported | NA |
| *Chamaecrista nictitans* | polyploid | polyploid | Senn HA (1938) Cytological Evidence of the Status of the Genus Chamaecrista Moench, Journal of the Arnold Arboretum, 19: 153-157. |
| *Cyclopia maculata* | polyploid | polyploid | Niemandt M, Roodt-Wilding R, Tobutt KR, Bester C (2018) Microsatellite marker applications in Cyclopia (Fabaceae) species, South African |

| | | | |
|---|---|---|---|
| | | | Journal of Botany, 116: 52-60. |
| *Dichilus lebeckioides* | polyploid | polyploid | Goldblatt P (1981) Chromosome Numbers in Legumes II, Missouri Botanical Garden, 68: 551-557. |
| *Dichilus strictus* | polyploid | polyploid | Goldblatt P (1981) Chromosome Numbers in Legumes II, Missouri Botanical Garden, 68: 551-557. |
| *Hesperolaburnum platycarpum* | polyploid | polyploid | Cusma-Velari T, Feoli-Chiapella L (2009) The so-called primitive genera of Genisteae (Fabaceae): systematic and phyletic considerations based on karyological data, Botanical Journal of the Linnean Society, 160: 232-248. |
| *Laburnum anagyroides* | polyploid | Not reported | NA |
| *Leucaena diversifolia* | polyploid | polyploid | Hughes CE, Bailey CD, Harris SA (2007) Divergent and reticulate species relationships in Leucaena (Fabaceae) inferred from multiple data sources: insights into polyploid origins and nrDNA polymorphism, American Journal of Botany, 89: 1057-1073. |
| *Leucaena greggii* | polyploid | diploid | Hughes CE, Bailey CD, Harris SA (2007) Divergent and reticulate species relationships in Leucaena (Fabaceae) inferred from multiple data sources: insights into polyploid origins and nrDNA polymorphism, American Journal of Botany, 89: 1057-1073. |
| *Leucaena involucrata* | polyploid | polyploid | Hughes CE, Bailey CD, Harris SA (2007) Divergent and reticulate species relationships in Leucaena (Fabaceae) inferred from multiple data sources: insights into polyploid origins and nrDNA polymorphism, American Journal of Botany, 89: 1057-1073. |
| *Leucaena multicapitula* | polyploid | diploid | Hughes CE, Bailey CD, Harris SA (2007) Divergent and reticulate species relationships in Leucaena (Fabaceae) |

| | | | |
|---|---|---|---|
| | | | inferred from multiple data sources: insights into polyploid origins and nrDNA polymorphism, American Journal of Botany, 89: 1057-1073. |
| *Leucaena pulverulenta* | polyploid | diploid | Hughes CE, Bailey CD, Harris SA (2007) Divergent and reticulate species relationships in Leucaena (Fabaceae) inferred from multiple data sources: insights into polyploid origins and nrDNA polymorphism, American Journal of Botany, 89: 1057-1073. |
| *Leucaena retusa* | polyploid | diploid | Hughes CE, Bailey CD, Harris SA (2007) Divergent and reticulate species relationships in Leucaena (Fabaceae) inferred from multiple data sources: insights into polyploid origins and nrDNA polymorphism, American Journal of Botany, 89: 1057-1073. |
| *Leucaena salvadorensis* | polyploid | diploid | Hughes CE, Bailey CD, Harris SA (2007) Divergent and reticulate species relationships in Leucaena (Fabaceae) inferred from multiple data sources: insights into polyploid origins and nrDNA polymorphism, American Journal of Botany, 89: 1057-1073. |
| *Leucaena shannonii* | polyploid | diploid | Hughes CE, Bailey CD, Harris SA (2007) Divergent and reticulate species relationships in Leucaena (Fabaceae) inferred from multiple data sources: insights into polyploid origins and nrDNA polymorphism, American Journal of Botany, 89: 1057-1073. |
| *Leucaena trichodes* | polyploid | diploid | Hughes CE, Bailey CD, Harris SA (2007) Divergent and reticulate species relationships in Leucaena (Fabaceae) inferred from multiple data sources: insights into polyploid origins and nrDNA polymorphism, American Journal of Botany, 89: 1057-1073. |
| *Neptunia oleracea* | polyploid | polyploid | Elias T (1974) The Genera of Mimosoideae (Leguminosae) in the |

| | | | |
|---|---|---|---|
| | | | Southeastern United States, The Arnold Arboretum of Harvard University, 55: 67-118. |
| *Neptunia plena* | polyploid | polyploid | Elias T (1974) The Genera of Mimosoideae (Leguminosae) in the Southeastern United States, The Arnold Arboretum of Harvard University, 55: 67-118. |
| *Peltophorum africanum* | polyploid | diploid | Van-Lume B, Souza G (2018) Cytomolecular analysis of species in the Peltophorum clade (Caesalpinioideae, Leguminosae), Brazilian Journal of Botany, 41: 385-392. |
| *Petteria ramentacea* | polyploid | polyploid/aneu ploid | Cusma-Velari T, Feoli-Chiapella L (2009) The so-called primitive genera of Genisteae (Fabaceae): systematic and phyletic considerations based on karyological data, Botanical Journal of the Linnean Society, 160: 232-248. |
| *Retama monosperma* | polyploid | polyploid | Benmiloud-Mahieddine R, Abirached-Darmency M, Brown SC, Kaid-Harche M, Siljak-Yakovlev S (2011) Genome size and cytogenetic characterization of three Algerian Retama species, Tree Genetics and Genomes, 7: 987-998. |
| *Retama raetam* | polyploid | polyploid | Benmiloud-Mahieddine R, Abirached-Darmency M, Brown SC, Kaid-Harche M, Siljak-Yakovlev S (2011) Genome size and cytogenetic characterization of three Algerian Retama species, Tree Genetics and Genomes, 7: 987-998. |
| *Retama sphaerocarpa* | polyploid | polyploid | Benmiloud-Mahieddine R, Abirached-Darmency M, Brown SC, Kaid-Harche M, Siljak-Yakovlev S (2011) Genome size and cytogenetic characterization of three Algerian Retama species, Tree Genetics and Genomes, 7: 987-998. |
| *Strongylodon macrobotrys* | polyploid | Not reported | NA |

| | | | |
|---|---|---|---|
| *Swartzia laevicarpa* | polyploid | polyploid strongly suggested | Pinto RB, de Freitas Mansano V, Torque BM, Forni-Martins ER (2015) Evidence for a conserved karyotype in Swartzia (Fabaceae, Papilionoideae): Implications for the taxonomy and evolutionary diversification of a species-rich netropical tree genus, Brittonia, 68: 93-101. |
| *Teramnus uncinatus* | polyploid | diploid | Lackey JA (1980) Chromosome Numbers in the Phaseoleae (Fabaceae:Faboideae) and their Relation to Taxonomy, American Journal of Botany, 67: 595-602. |
| *Virgilia oroboides* | polyploid | Not reported | NA |

**Supplemental Table 4.** Diploid species identified in our dataset. We looked up a randomly selected 62 species in the literature to confirm if they were diploid or whether we should correct our dataset and change certain species to polyploid.

| Species | Ploidy level in our dataset | Ploidy level in the literature | Reference |
|---|---|---|---|
| *Bauhinia petersiana* | diploid | likely diploid | Kumar A, Raju DT (1967) Structure and Behaviour of Chromosomes in Bauhinia and Allied Genera, Cytologia, 33: 411-426. |
| *Pueraria phaseoloides* | diploid | Not reported | NA |
| *Vigna marina* | diploid | diploid | Tomooka N et al (2014) Evolution, domestication and neo-domestication of the genus Vigna, Plant Genetic Resources, 12: S168-S171 |
| *Crotalaria comosa* | diploid | Not reported | NA |
| *Amorpha canescens* | diploid | maybe polyploid | Delaney JT, Baack EJ (2012) Intraspecific Chromosome Number Variation and Prairie Restoration - A Case Study in Northeast Iowa, USA, Restoration Ecology, 20: 576-583. |
| *Vicia pannonica* | diploid | diploid | Yeater KM, Bollero GA, Bullock DG, Rayburn AL (2004) Flow cytometric analysis for ploidy level differentitation of 45 hariy vetch accessions, Ann. Appl. Biol. 145: 123-127 |
| *Trifolium pratense* | diploid | naturally diploid | Jing S, Kryger P, Boelt B (2021) Different pollination approaches to compare the seed set of diploid and tetraploid red clover Trifolium pratense L. Nordic Journal of Botany, 39: e03006. |
| *Crotalaria incana* | diploid | diploid | Gupta PK, Gupta R (1977) Pollen morphology in diploid species of Crotalaria L, Proc. Indian Acad. Sci. 88: 49-56. |
| *Medicago suffruticosa* | diploid | likely diploid | Gillies CB (1972) Pachytene chromosomes of perennial Medicago species, Hereditas, 72: 303- |

| | | | |
|---|---|---|---|
| | | | 310. |
| *Sophora chrysophylla* | diploid | diploid | Sousa SM, Rudd VE (1993) Revision del Genero Styphnolobium (Leguminosae: Papilionoideae: Sophoreae). Annals. Of the Misouri Botanical Garden, 80: 270-283. |
| *Gymnocladus dioica* | diploid | likely diploid | Schnabel A, Wendel JF (1998) Cladistic biogeography of Gleditsia (Leguminosae) based on ndhF and rpl16 chloroplast gene sequences, American Journal of Botany, 85: 1753-1765. |
| *Dalbergia nigra* | diploid | likely diploid | Silva Júnior AL (2020) Evaluation of diversity and genetic structure as strategies for conservation of natural populations of Dalbergia nigra (Vell.) Allemão ex Benth., CERNE, 26: 435-443). |
| *Sindora supa* | diploid | diploid | Robertson KR, Lee YT (1976) The Genera of Caesalpinioideae (Leguminosae) in the Southeastern United States, The Arnold Arboretum of Harvard University, 57: 1-53. |
| *Mimosa polycarpa* | diploid | diploid | Dahmer N et al (2010) Chromosome numbers in the genus Mimosa L.: cytotaxonomic and evolutionary implications, Plant Systematics and Evolution, 291: 211-220. |
| *Daviesia ulicifolia* | diploid | Not reported | NA |
| *Coronilla juncea* | diploid | diploid | Dušková J, Sovová M, Žáčková P, Spurná V (1987) Tissue Culture of Crownvetch (Coronilla varia L.) and the Production of Cardenolide-Like Substances in vitro, Biologia Plantarum, 29: 258-264. |
| *Hypocalyptus coluteoides* | diploid | Not reported | NA |
| *Lathyrus cicera* | diploid | diploid | Unal F (1993) Genomic and chromosomal evolution in Lathyrus, The University of Manchester (United Kingdom). |
| *Medicago* | diploid | diploid | Quiros CF, Ostafichuk L (1983) Allozymes and |

| | | | |
|---|---|---|---|
| *turbinata* | | | genetic variability in Medicago turbinata, M. truncatula, and their hybrids, Canadian Journal of Genetics and Cytology, 25: 286-291. |
| *Lespedeza angustifolia* | diploid | diploid | Xu B, Zeng, XM, Gao XF, Jin DP, Zhang LB (2017) ITS non-concerted evolution and rampant hybridization in the legume genus Lespedeza (Fabaceae), Scientific Reports, 7: 40057 |
| *Vigna mungo* | diploid | naturally diploid | Singh MN, Singh SK (2005) Study of induced amphidiploid derivatives of Vigna radiata x Vigna mungo, Indian J. Genet., 66: 245-246. |
| *Sesbania herbacea* | diploid | diploid | Kumar S, Friebe B, Gill BS (2014) Physical localization of rRNA genes by fluorescence in situ hybridization (FISH) and analysis of spacer length variants of 45S rRNA (slvs) genes in some species of genus Sesbania, Plant Systematics and Evolution, 300: 1793-1802. |
| *Brya ebenus* | diploid | Not reported | NA |
| *Lathyrus japonicus* | diploid | diploid | Barrington DS, Schmitz SA (2013) Quaternary divergence and holocene secondary contact via the northwest passage in the circumpolar Lathyrus japonicus (Leguminosae), Rhodora, 115: 133-157. |
| *Platylobium formosum* | diploid | Not reported | NA |
| *Daviesia acicularis* | diploid | Not reported | NA |
| *Dumasia villosa* | diploid | diploid | Kumar PS, Hymowitz TH (1989) Where are the diploid (2n=2x=20) genome donors of Glycine Willd. (Leguminosae, Papilionoideae)?, Euphytica, 40:221-226. |
| *Inocarpus fagifer* | diploid | Not reported | NA |
| *Phaseolus vulgaris* | diploid | diploid | McClean PE, Lavin M, Gepts P, Jackson SA (2008) Phaseolus vulgaris: a diploid model for |

| | | | |
|---|---|---|---|
| | | | soybean, Genetics and genomics of soybean, Springer, New York, NY, 55-76. |
| *Trifolium microcephalum* | diploid | diploid | Vozárová R, Macková E, Vik D, Řepková J (2021) Variation in Ribosomal DNA in the genus Trifolium (Fabaceae), Plants, 10: 1771. |
| *Indigastrum argyroides* | diploid | diploid | Frahm-Leliveld JA (1966) Cytotaxonomic notes on the genera Indigofera L. and Cyamopsis DC., Genetica, 37: 403-426. |
| *Stryphnodendron adstringens* | diploid | Not reported | NA |
| *Acacia ampliceps* | diploid | diploid | Ismail, M., Ahmad, A., Nadeem, M., Javed, M. A., Khan, S. H., Khawaish, I., Qamer, S. (2020). Development of DNA barcodes for selected Acacia species by using rbcL and matK DNA markers. *Saudi Journal of Biological Sciences*, *27:* 3735-3742. |
| *Centrolobium tomentosum* | diploid | diploid | Cordeiro, E. M., Macrini, C. M., Sujii, P. S., Schwarcz, K. D., Pinheiro, J. B., Rodrigues, R. R., ... & Zucchi, M. I. (2019). Diversity, genetic structure, and population genomics of the tropical tree Centrolobium tomentosum in remnant and restored Atlantic forests. *Conservation Genetics*, *20*(5), 1073-1085. |
| *Trifolium squamosum* | diploid | diploid | Vozárová, R., Macková, E., Vlk, D., & Řepková, J. (2021). Variation in ribosomal DNA in the genus Trifolium (Fabaceae). Plants, 10(9), 1771. |
| *Oxytropis splendens* | diploid | diploid | Archambault, A., & Strömvik, M. V. (2012). Evolutionary relationships in Oxytropis species, as estimated from the nuclear ribosomal internal transcribed spacer (ITS) sequences point to multiple expansions into the Arctic. *Botany*, *90*(8), 770-779. |
| *Lespedeza floribunda* | diploid | diploid | Young, J. O. (1940). Cytological investigations in Desmodium and Lespedeza. *Botanical Gazette*, *101*(4), 839-850 |

| | | | |
|---|---|---|---|
| *Medicago noeana* | diploid | diploid | Mehregan, I., Rahiminejad, M. R., & Azizian, D. (2002). A taxonomic revision of the genus Medicago L.(Fabaceae) in Iran. *Iranian Journal Botany*, *9*(2), 207-221. |
| *Cicer arietinum* | diploid | diploid | Thomas, P. T., & Revell, S. H. (1946). Secondary association and heterochromatic attraction: I. Cicer arietinum. *Annals of Botany*, *10*(38), 159-164 |
| *Piscidia piscipula* | diploid | Not reported | NA |
| *Calliandra houstoniana* | diploid | Not reported | NA |
| *Dalea candida* | diploid | likely diploid | Moncada, K. M., Ehlke, N. J., Muehlbauer, G. J., Sheaffer, C. C., Wyse, D. L., & DeHaan, L. R. (2007). Genetic variation in three native plant species across the state of Minnesota. *Crop Science*, *47*(6), 2379-2389. |
| *Neptunia pubescens* | diploid | polyploid | Silva, J. S., Carvalho, M. S., Santos, G. S., Braga, F. T., Gomes de Andrade, M. J., & de Freitas Mansano, V. (2020). Neptunia windleriana: A New Polyploid Species of Neptunia (Leguminosae) from Brazil Recognized by Anatomy, Morphology and Cytogenetics. *Systematic Botany*, *45*(3), 483-494. |
| *Dalbergia assamica* | diploid | diploid | Hiremath, S. C., & Nagasampige, M. H. (2004). Genome size variation and evolution in some species of Dalbergia Linn. f.(Fabaceae). *Caryologia*, *57*(4), 367-372. |
| *Trifolium squarrosum* | diploid | diploid | Falistocco, E., Marconi, G., & Falcinelli, M. (2013). Comparative cytogenetic study on Trifolium subterraneum (2 n= 16) and Trifolium israeliticum (2 n= 12). *Genome*, *56*(6), 307-313. |
| *Sophora prostrata* | diploid | Not reported | NA |

| | | | |
|---|---|---|---|
| *Vicia unijuga* | diploid | diploid | Han, Y., & Wang, H. Y. (2010). Genetic diversity and phylogenetic relationships of two closely related northeast China Vicia species revealed with RAPD and ISSR markers. *Biochemical genetics*, *48*(5-6), 385-401 |
| *Acacia caffra* | diploid | diploid | Van Drunen, W. (2018). The association between polyploidy and clonal reproduction in the angiosperms (Doctoral dissertation). |
| *Hippocrepis emerus* | diploid | diploid | Lonn, M., & Prentice, H. C. (1995). The structure of allozyme and leaf shape variation in isolated, range-margin populations of the shrub Hippocrepis emerus (Leguminosae). *Ecography*, *18*(3), 276-285 |
| *Albizia guachapele* | diploid | likely diploid | ARCE, M. D. L. R. (1992). New chromosome counts in neotropical Albizia, Havardia and Pithecellobium, and a new combination for Albizia (Leguminosae-Mimosoideae-Ingeae). *Botanical Journal of the Linnean Society*, *108*(3), 269-274 |
| *Trifolium rueppellianum* | diploid | diploid | Ellison, N. W., Liston, A., Steiner, J. J., Williams, W. M., & Taylor, N. L. (2006). Molecular phylogenetics of the clover genus (Trifolium—Leguminosae). *Molecular phylogenetics and evolution*, *39*(3), 688-705 |
| *Lespedeza stuevei* | diploid | diploid | Xu, B., Zeng, X. M., Gao, X. F., Jin, D. P., & Zhang, L. B. (2017). ITS non-concerted evolution and rampant hybridization in the legume genus Lespedeza (Fabaceae). *Scientific Reports*, *7*(1), 1-15. |
| *Aeschynomene pratensis* | diploid | diploid | Chaintreuil, C., Perrier, X., Martin, G., Fardoux, J., Lewis, G. P., Brottier, L., ... & Arrighi, J. F. (2018). Naturally occurring variations in the nod-independent model legume Aeschynomene evenia and relatives: a resource for nodulation genetics. *BMC plant biology*, *18*(1), 1-15. |
| *Hedysarum* | diploid | likely diploid | Nuzhdina, N. S., Bondar, A. A., & Dorogina, O. |

| | | | |
|---|---|---|---|
| *vicioides* | | | V. (2018). New Data on Taxonomic and Geographic Distribution of the trn L UAA Intron Deletion of Chloroplast DNA in Hedysarum L.(Fabaceae L.). *Russian Journal of Genetics*, *54*(11), 1282-1292 |
| *Lespedeza juncea* | diploid | diploid | Xu, B., Zeng, X. M., Gao, X. F., Jin, D. P., & Zhang, L. B. (2017). ITS non-concerted evolution and rampant hybridization in the legume genus Lespedeza (Fabaceae). Scientific Reports, 7(1), 1-15. |
| *Desmodium barbatum* | diploid | diploid | Adelanwa MA, Husaini SWA (2014) Cytomorphological studies of the tribe Desmodieae found in northern Nigeria, International Journal of Agriculture Innovations and Research, 2: 2319-1473. |
| *Acmispon parviflorus* | diploid | Not reported | NA |
| *Indigofera demissa* | diploid | likely diploid | Frahm-Leliveld, J. A. (1962). Further Observations on Chromosomes in the Genus Indigofera L. *Acta botanica neerlandica*, *11*(2), 201-208. |
| *Securigera securidaca* | diploid | Not reported | NA |
| *Lotus edulis* | diploid | diploid | Gauthier, P., Lumaret, R., & Bedecarrats, A. (1997). Chloroplast-DNA variation in the genus Lotus (Fabaceae) and further evidence regarding the maternal parentage of Lotus corniculatus L. *Theoretical and Applied Genetics*, *95*(4), 629-636 |
| *Albizia procera* | diploid | Not reported | NA |
| *Vicia parviflora* | diploid | Likely diploid | Han, Y., & Wang, H. Y. (2010). Genetic diversity and phylogenetic relationships of two closely related northeast China Vicia species revealed with RAPD and ISSR markers. *Biochemical genetics*, *48*(5-6), 385-401 |

**Supplemental Table 5.** PGLS estimates and results for effects of ploidy and symbiotic status on the number of introduced ranges in legumes. Ploidy is estimated from combined genus and subfamily-level base chromosome numbers. Italicized factors are significant at $p < 0.05$ and p values reported here are the results of type III ANOVAs. Bolded factors highlight significant interaction effects on legume introductions. Ploidy was coded as 0=diploid and 1=polyploid while symbiotic status was coded as 0=non-symbiotic and 1=symbiotic in the model. Therefore the intercept represents non-symbiotic diploids. Specialists were coded as 1 and generalists as 0 in the model.

| Factor | Estimate | s.e. | d.f. | Wald $\chi^2$ | p value |
|---|---|---|---|---|---|
| **Symbiosis status, d.f. = 811, $\lambda$ = 0.0431** | | | | | |
| Ploidy (polyploid) | 0.124 | 0.094 | 1 | 1.726 | 0.1889 |
| Symbiosis (symbiotic) | -0.006 | 0.057 | 1 | 0.009 | 0.9252 |
| *Total native area* | *-0.042* | *0.012* | *1* | *12.475* | *0.0004* |
| *Human uses* | *0.188* | *0.007* | *1* | *748.492* | *< 0.0001* |
| *Absolute latitude* | *-0.002* | *0.001* | *1* | *5.560* | *0.0180* |
| *Annual* | *0.073* | *0.033* | *1* | *4.753* | *0.0292* |
| Ploidy*Symbiosis | -0.103 | 0.099 | 1 | 1.086 | 0.2973 |
| **Specialist status, d.f. = 130, $\lambda$ = - 0.1992** | | | | | |
| *Ploidy (polyploid)* | *0.479* | *0.172* | *1* | *7.778* | *0.0053* |
| *Specificity (specialist)* | *0.168* | *0.087* | *1* | *3.706* | *0.0542* |
| *Total native area* | *-0.116* | *0.024* | *1* | *23.923* | *< 0.0001* |
| *Human uses* | *0.173* | *0.015* | *1* | *125.239* | *< 0.0001* |
| *Absolute latitude* | *-0.012* | *0.002* | *1* | *26.422* | *< 0.0001* |
| Annual | 0.007 | 0.098 | 1 | 0.006 | 0.9391 |
| ***Ploidy*Specificity*** | ***-0.530*** | ***0.192*** | ***1*** | ***7.597*** | ***0.0058*** |

**Supplemental Table 6.** Results of PGLS and glm models testing the effect of ploidy on introduction success in two categories of the data: symbiotic and non-symbiotic species. Italicized factors are significant at $p < 0.05$ and reported p values reported here are the results of type II ANOVAs. Ploidy was coded as 0=diploid and 1=polyploid.

| **Factor** | **Estimate** | **s.e.** | **d.f.** | **Wald $\chi^2$** | **p value** |
|---|---|---|---|---|---|
| **Non-symbiotic species, PGLS, d.f. = 66, $\lambda$ = 0.3552** | | | | | |
| *Ploidy (polyploid)* | *0.256* | *0.115* | *1* | *4.564* | *0.0327* |
| Total native area | 0.110 | 0.063 | 1 | 3.075 | 0.0795 |
| *Human uses* | *0.170* | *0.022* | *1* | *59.450* | *< 0.0001* |
| Absolute latitude | 0.007 | 0.004 | 1 | 2.845 | 0.0916 |
| Annual | 0.125 | 0.417 | 1 | 0.060 | 0.7647 |
| **Symbiotic species, PGLS, d.f. = 745, $\lambda$ = 0.0401** | | | | | |
| Ploidy (polyploid) | 0.018 | 0.029 | 1 | 0.395 | 0.5298 |
| *Total native area* | *-0.052* | *0.012* | *1* | *18.752* | *0.0062* |
| *Human uses* | *0.195* | *0.007* | *1* | *729.775* | *< 0.0001* |
| *Absolute latitude* | *-0.003* | *0.001* | *1* | *7.491* | *0.0062* |
| *Annual* | *0.088* | *0.033* | *1* | *7.258* | *0.0070* |
| **Symbiotic species, quasi-poisson, d.f. = 744** | | | | | |
| Ploidy (polyploid) | -0.071 | 0.123 | 1 | 0.440 | 0.5049 |
| *Total native area* | *-0.144* | *0.043* | *1* | *17.640* | *< 0.0001* |
| *Human uses* | *0.438* | *0.019* | *1* | *542.870* | *< 0.0001* |
| *Absolute latitude* | *-0.012* | *0.004* | *1* | *14.280* | *0.0002* |
| *Annual* | *0.413* | *0.138* | *1* | *12.250* | *0.0005* |
